# High-throughput single-cell TCR - pMHC dissociation rate measurements performed by an autonomous microfluidic cellular processing unit

**DOI:** 10.1101/2021.06.30.450499

**Authors:** Fabien Jammes, Julien Schmidt, George Coukos, Sebastian J. Maerkl

## Abstract

We developed an integrated microfluidic cellular processing unit (mCPU) capable of autonomously isolating single cells, perform, measure, and on-the-fly analyze cell-surface dissociation rates, followed by recovery of selected cells. We performed proof-of-concept, high-throughput single-cell experiments characterizing pMHC-TCR interactions on live CD8+ T cells. The mCPU platform analyzed TCR-pMHC dissociation rates with a throughput of 50 cells per hour and hundreds of cells per run, and we demonstrate that cells can be selected, enriched, and easily recovered from the device.

---

Microfluidics, or “lab-on-a-chip” technology has the potential to autonomously perform complex laboratory methods. In mammalian cell applications, microfluidic devices automated tasks such as long-term cell culture [1], transfection [2], cell-secretion analysis [3], chemical cytometry [4, 5], cell sorting [6], and many others. In most if not all applications to date, microfluidic devices are used to scale-down and automate experimental processes, but data processing and analysis are generally performed off-line after experiments have been completed.

While microfluidic platforms have been applied to single-cell applications, including peptide - major histocompatibility complexes (pMHC) and T cell receptors (TCR) interactions [7, 8], no platforms have so far enabled a closed-loop selection system with spatial and temporal resolution at the single-cell level. Droplet microfluidic technologies enabled high-throughput screening and sorting of immune cells [9, 10] but rely on a single time point measurements for analysis and decision making, which limits the applicability to static or slow biological processes. On the other hand, microfluidic platforms capable of analysing single cells in response to stimuli have been described but have achieved limited throughput analyzing tens of cells per experiment and do not enable specific cell recovery [11]. The state-of-the-art for single cell TCR - pMHC dissociation rate measurements remain low-throughput, non-automated approaches, which also lack the ability to selectively recover cells [12]. Stockslager et al. developed a microfluidic approach using surface immobilized pMHC combined with flowing T cells over the pMHC surface. A reduction in cell-speed indicates that cells interact with the surface immobilized pMHCs [8]. A few dozen cells were analyzed with this approach, but it is not clear whether the method can differentiate differences in pMHC - TCR affinity and doesn’t provide a direct measurement of pMHC - TCR dissociation rates. To the best of our knowledge no microfluidic approaches have been described that conduct dissociation rate measurements of pMHC - TCR complexes on single cells.

Here we describe a fully-autonomous microfluidic cellular processing unit (mCPU) that performs single-cell surface marker dissociation measurements, and runs all necessary fluid handling, microscope, imaging, image processing, and data analysis steps on-the-fly, creating a closed-loop system (Fig. 1 and Supplementary Fig. S1). Machine vision and image analysis is used to feedback and control device operation allowing the device to perform complex fluidic processing steps autonomously. We applied this platform to a proof-of-concept, high-throughput measurement determining the dissociation rates between peptide -major histocompatibility complexes and T cell receptors on hundreds of cells, and demonstrate that the platform can automatically select cells based on the measured dissociation rate, and recover these cells from the device.

**Figure 1:**
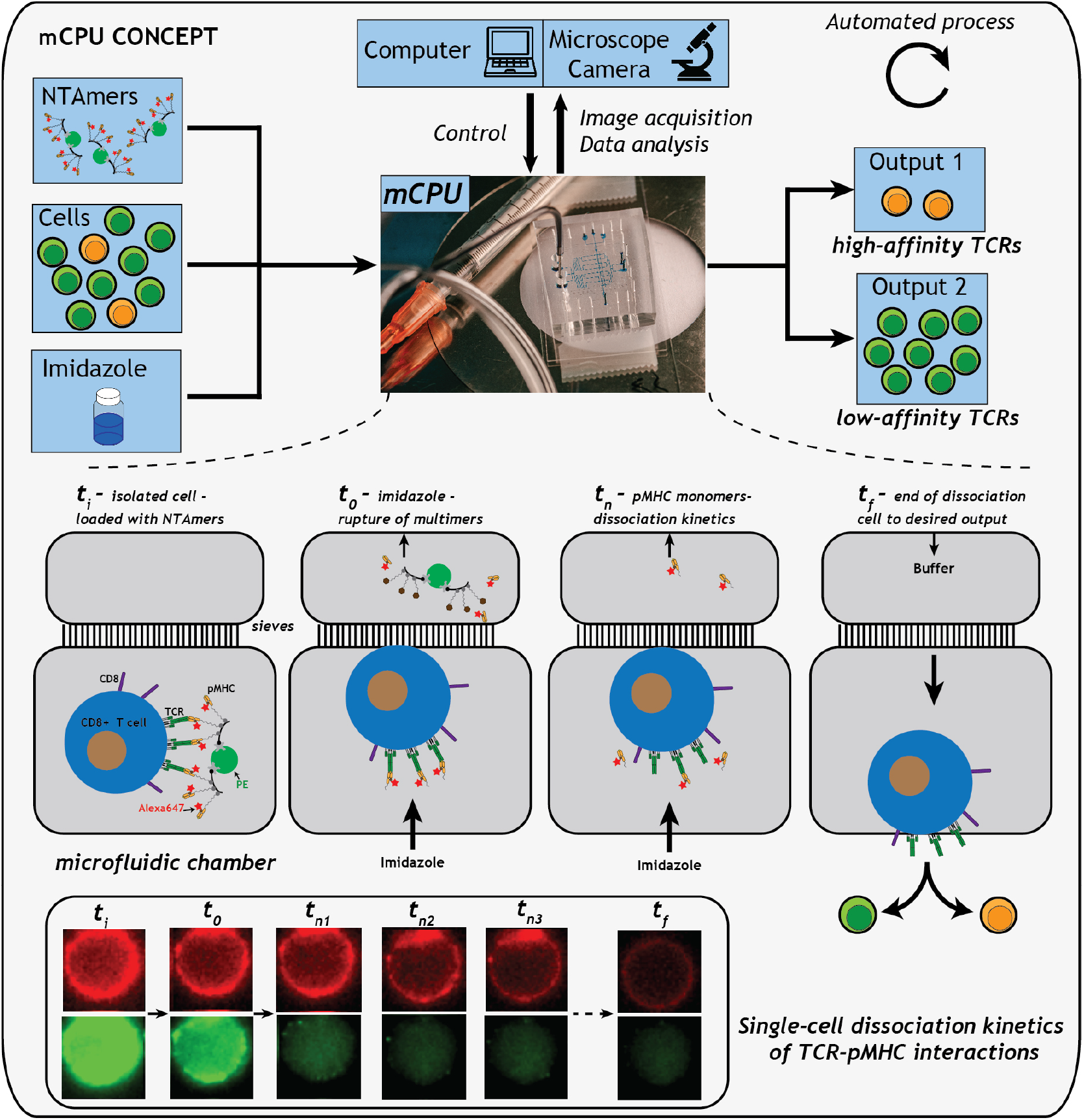
mCPU concept & design. A control chart describing the mCPU platform. A microfluidic device forms the core of the mCPU. The microfluidic device is composed of individual microfluidic chambers capable of single-cell isolation and analysis of TCR-pMHC dissociation over time. We used pMHC NTAmers [16] for dissociation measurements of cell-surface TCR-pMHC interactions. Cells stained with pMHC NTAmers are isolated in individual chambers, followed by a rapid buffer exchange introducing imidazole which dissociates the NTAmer, leaving pMHC monomers bound to the cell surface which dissociate over time according to the pMHC-TCR affinity. Time-lapse imaging allows determination of the pMHC dissociation rate. This enables automated separation and selection of low- and high-affinity TCRs from a mixed population.

We demonstrate that the mCPU can measure the dissociation rate of the interaction between TCRs on CD8+ T cells and pMHCs. This affinity has been suggested to be an important parameter and to correlate with T cell function [12, 13, 14], although the importance of mechanical force to the dissociation rate remains controversial [15]. The mCPU platform achieves sensitive, dynamic, single-cell analysis and high-throughput by enabling autonomous and closed-loop temporal analysis of single cells, while enabling recovery and enrichment of selected cells.

## Results

The mCPU design consists of two fluidic layers, 106 micromechanical valves, and 8 cell isolation chambers in which cells are isolated and analyzed (Fig. 2a and Supplementary Fig. S2). Buffer conditions can be changed in each chamber by fast flow exchange without loss of cells by incorporating small sieves and fluidic bypasses in the chamber design (Fig. 2b). Complete automation of the experimental process was achieved via a MATLAB program that controls the microfluidic device, camera, and microscope (Fig. 2c). To achieve autonomous single-cell experiments, the mCPU platform performs sequential experimental rounds consisting of cell loading and isolation, followed by automatic cell detection in the chambers (Fig. 2d). Buffer exchange is triggered in chambers identified to contain a cell, followed by data acquisition through time-lapse imaging. An elaborate analytical pipeline performs image processing, data extraction, and curve-fitting for real-time extraction of single-cell surface marker dissociation rates (Supplementary Figures S3 and S4). Cells above or below a particular dissociation rate threshold can be selected for recovery. The system then automatically triggers the analysis process in the next cell-containing chamber, or, if no more chambers contain a cell, triggers a new round of cell loading starting the next experimental cycle (Fig. 2d).

**Figure 2:**
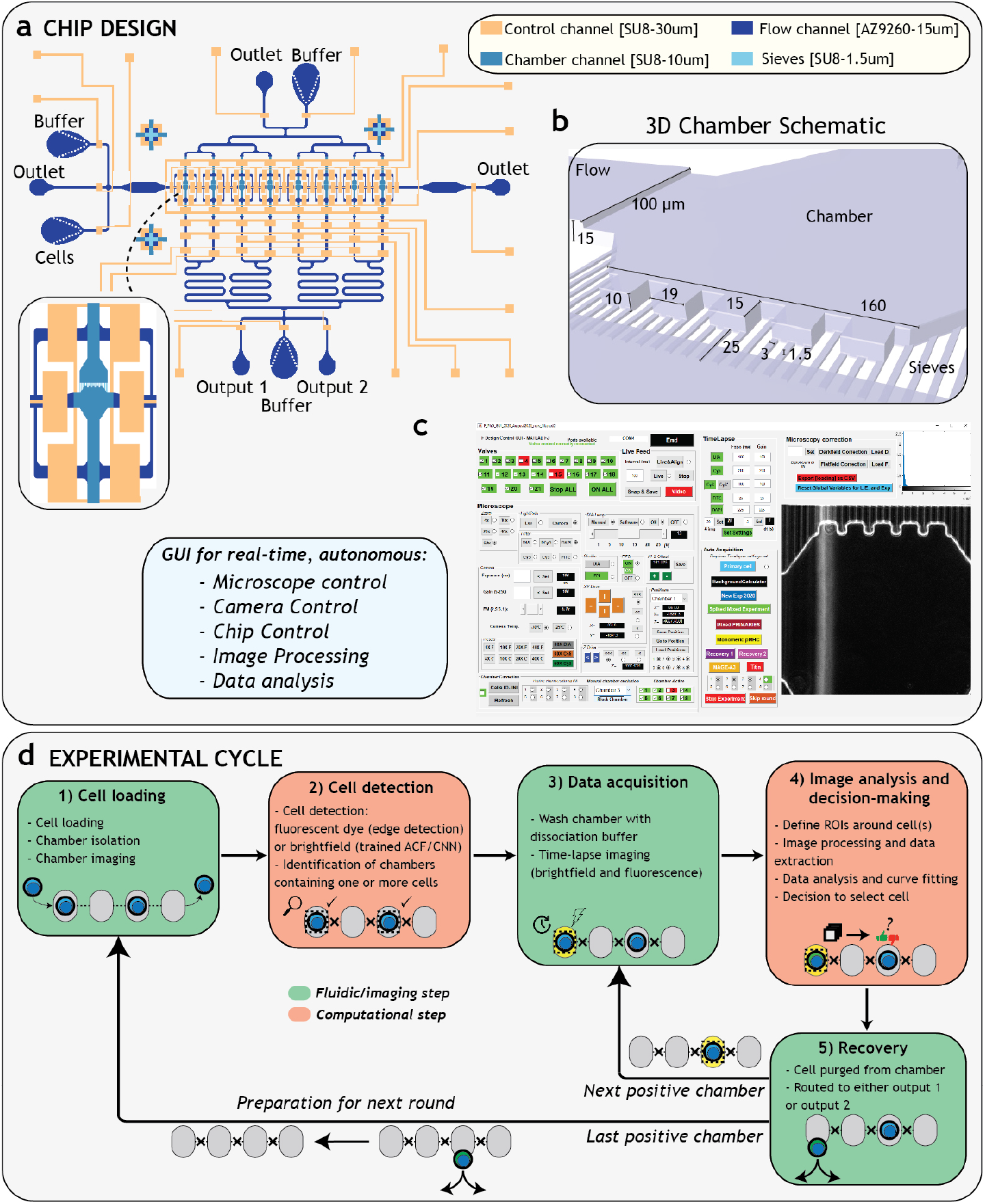
mCPU concept & design. **a** The microfluidic device design showing the two-layer channel network consisting of a control (orange) and flow (blue) layer. **b** 3D chamber schematic showing chamber design details. **c** Automation is achieved through a custom MATLAB GUI controlling the chip, camera, and microscope, and performs all image processing and data analysis steps in real-time. **d** The mCPU platform performs sequential experimental rounds that are divided into five discrete steps.

Here we used NTAmers which are pMHC multimers that can be rapidly dissociated by imidazole (Fig. 1) [16]. To perform dissociation rate measurements of T-cell receptor - pMHC interactions, TCR-transduced CD8+ SUP-T1 cells were pre-stained with NTAmer-pMHC multimers followed by on-chip analysis. Cell containing chambers were identified automatically using either brightfield images or via fluorescence using live-cell stains (Fig. 3a,b and Supplementary Fig. S5). In 6 experimental runs the mCPU achieved an average throughput of ∼50 cells per hour and ran stably and continuously for at least 6 hours (Fig. 3c). Acquisition of dissociation kinetics was performed by rapid (≤5 seconds) buffer exchange introducing an imidazole containing buffer in the chamber and acquired images over a period of 60 to 90 seconds with a 3 second interval (Fig. 3d). Image analysis allowed for automatic determination of the switch from multimeric to monomeric forms by detecting the sudden decrease in PE signal (Fig. 3e). Each event was filtered based on multiple parameters described in the methods section (Supplementary Figures S6 and S7). Monomeric pMHC - TCR dissociation (Alexa647) was then fitted to a one-phase exponential decay model (Supplementary Fig. S8). Finally, the cell (or cells) in the chamber was directed to one of two chip outputs depending on whether it met a minimal dissociation rate threshold (Fig. 3f,g).

**Figure 3:**
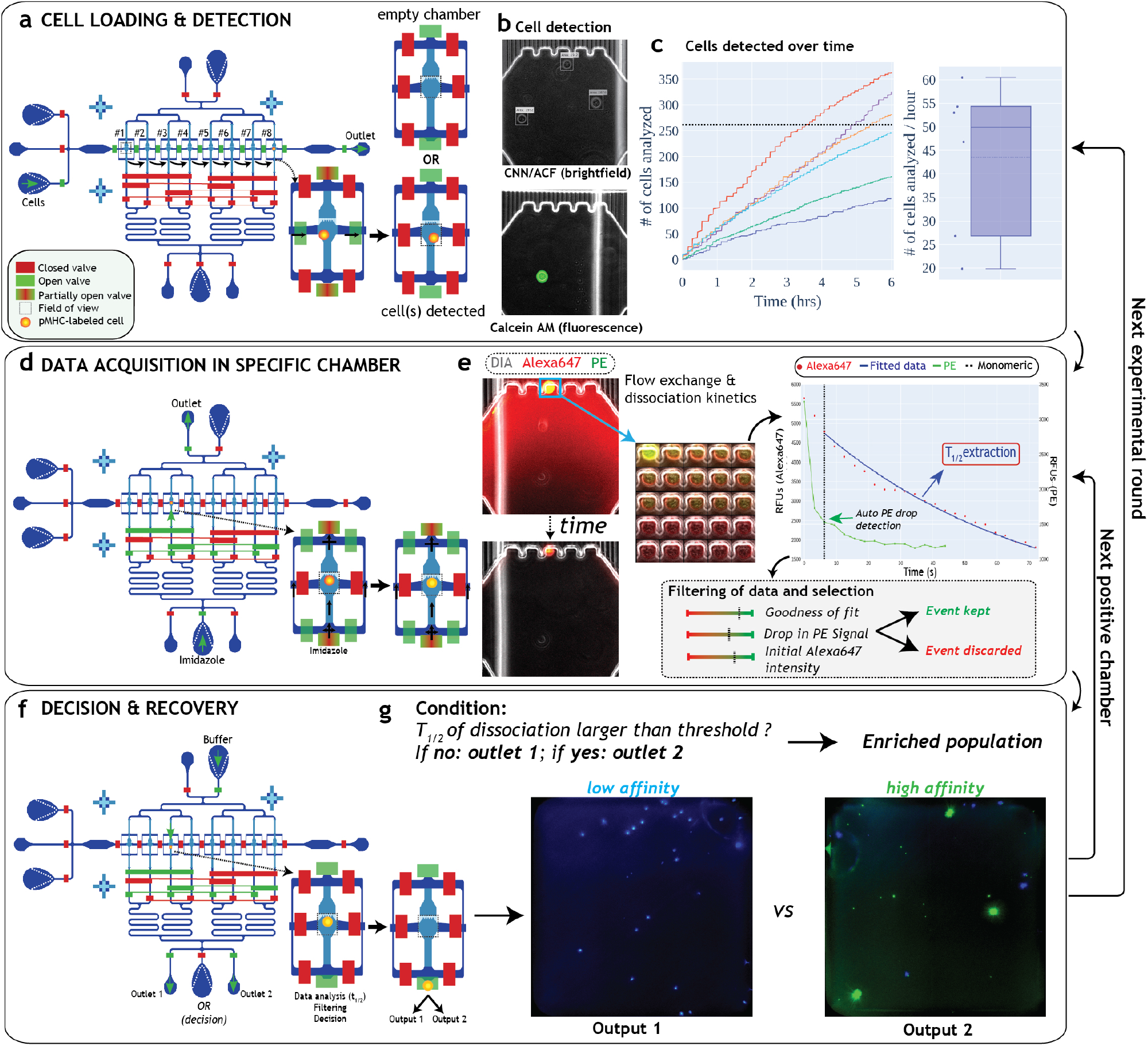
mCPU single-cell analysis. **a-g** The mCPU performs sequential microfluidic processing steps: cell-loading and detection, buffer exchange and single-cell analysis, and cell selection and recovery **a** Cells are loaded and the eight chambers are isolated and imaged to identify chambers containing one or more cells. **b** Cell detection is performed either in brightfield or fluorescence. **c** The cumulative number of cells recorded and analyzed in all chambers per round of cell loading, showing 6 separate experiments. The average analysis rate was ∼50 cells per hour. **d** Chambers containing one or more cells are then imaged via fluorescent time-lapse imaging and washed with imidazole buffer leading to NTAmer dissocation. **e** Time-lapse acquisition is performed on two fluorescent channels: PE (NTAmer dissociation), and Alexa647 (pMHC - TCR dissociation). Image processing, data analysis, and curve fitting is performed to extract TCR-pMHC dissociation half-lives. **f-g** Filtered data are then compared with user-defined thresholds for cell selection and routing to a specific outlet. The process is then repeated in the next cell containing chamber or a new round of cell-loading is triggered.

We focused on characterizing a wild-type (WT), low-affinity patient-derived TCR specific for the NY-ESO-1 tumor antigen and a higher affinity variant (DM*β*) that was previously generated [17]. We bench-marked our system by measuring both the WT and DM*β* variants, as they cover the physiologically relevant dissociation rates ranging from high for WT (short half-lives) to low for DM*β* (long half-lives) [18].

In six mCPU runs a total of 908 cells were analyzed, 447 WT and 461 DM*β* expressing cells. We obtained average half-lives of 29.1s and 80.5s for WT and DM*β*, respectively (Fig. 4a; Supplementary Figures S9 & S10). These half-lives are similar to those obtained by a clonal population-level FACS measurement requiring at least 200’000 cells (27.1s and 75.8s for WT and DM*β* respectively, Supplementary Fig. S11) and a previous FACS analysis (17s and 89.7s for WT and DM*β*, respectively [19]). The results were repeatable between experiments based on the observed half-life averages (Fig. 4b). We measured coefficients of variation of 55% and 56% for WT and DMB, based on the analysis of 447 and 461 cells, respectively, across 3 technical repeats. By comparison, previous measurements based on the analysis of 51-60 cells resulted in coefficients of variation of 23-29% [12]. Whether this increased variance is primarily of biological or technical origin remains, or intrinsic to the cell type used in this study remains to be determined. In this series of experiments 82% of chambers contained a single cell, 14% contained 2 cells, and 4% contained 3 or more cells identified pre-filtering (Fig. 4c). These experiments demonstrate that the mCPU is capable of autonomously performing large-scale, single-cell surface marker dissociation measurements, which should find uses in characterizing T cell, tumor-infiltrating lymphocyte (TIL), or B cell samples, and synthetically generated surface receptor libraries.

**Figure 4:**
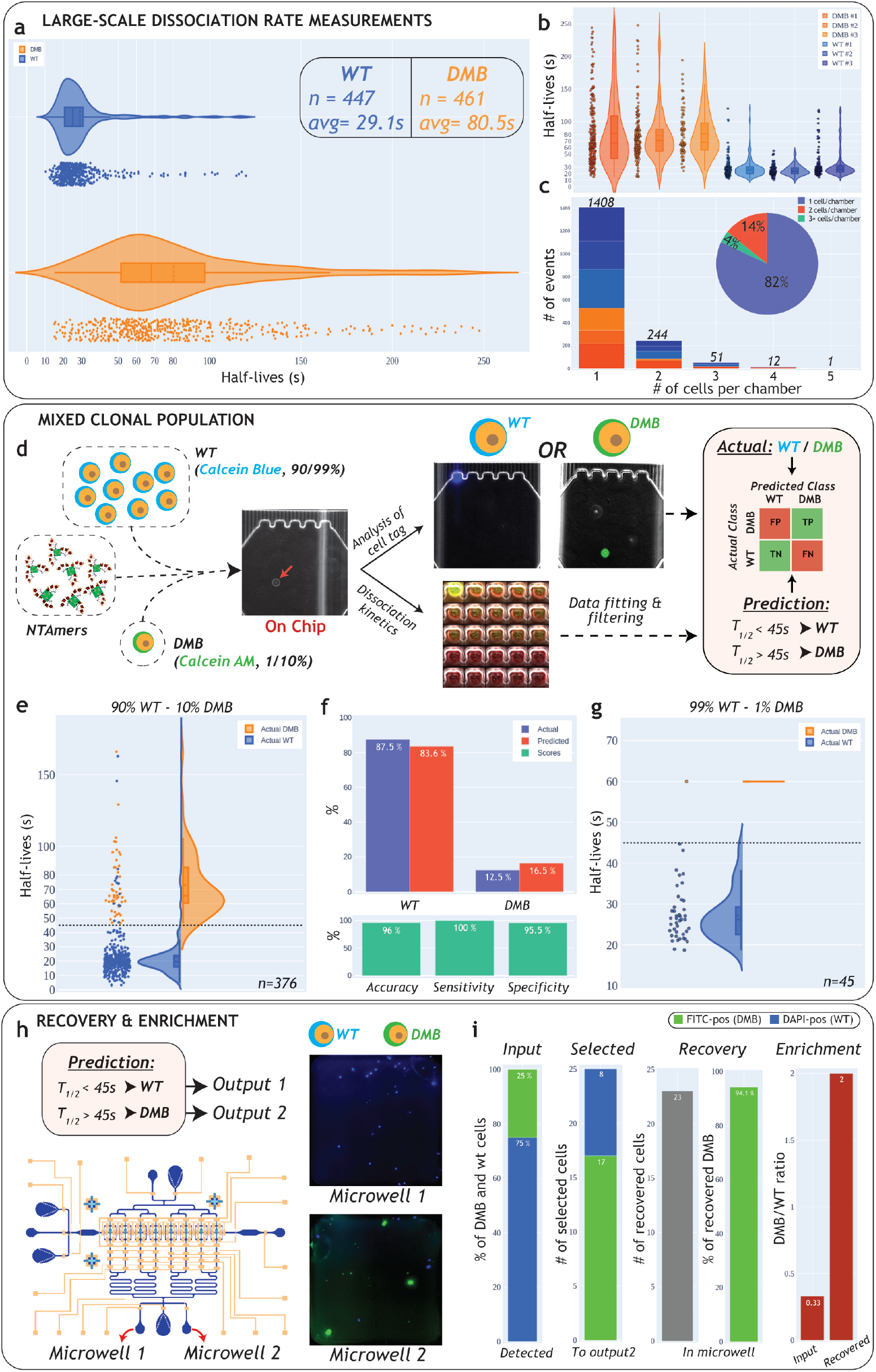
Single-cell TCR-pMHC dissociation measurements. **a** Large-scale single-cell dissociation rate measurements for WT and DM*β* expressing cells (aggregate of 3 independent experiments for each clone). **b** Half-live distributions for each experiment. **c** Distribution of cells detected per chamber. **d** Mixed population experiments. The actual WT to DM*β* ratio was determined on-chip via Calcein Blue for WT and Calcein AM for DM*β*, followed by dissociation rate measurements. **e** 90% WT to 10% DM*β* experiment. Measured half-lives for each cell. Data point color indicates cell-type as determined by calcein staining. A half-life threshold of 45s was used to differentiate between high and low affinity cells. **f** WT to DM*β* ratio as determined by calcein (actual) and as determined by using half-life measurements with a 45s half-life threshold (predicted). Accuracy, sensitivity and specificity were determined based on these ratios. **g** Experiment with higher dilution of high-affinity cells (99%WT and 1%DM*β*) performed to identify a single-high affinity clone. **h** A recovery and enrichment experiment was performed by routing selected cells to a specific chip outlet. Cells were then transferred into microwells for counting. **i** Recovery and enrichment ratios were determined by comparing on-chip input with counted cells in microwells

**Figure 5:**
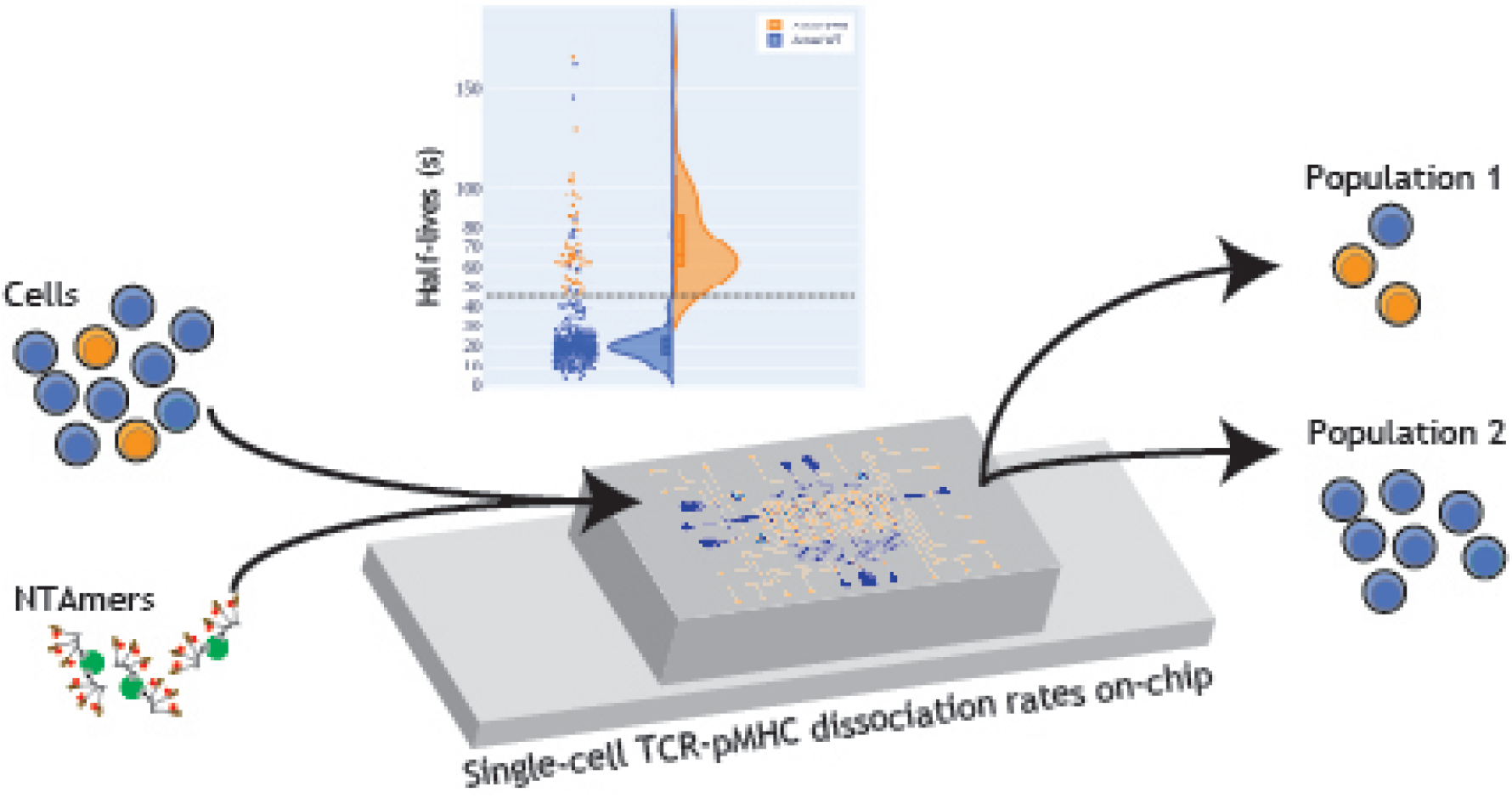
for TOC.

The mCPU integrates cell selection and recovery capabilities to extract cells of interest from the mCPU for downstream processing. To test the mCPU’s capability of identifying and enriching rare cells from a mixed population we generated a mixture of WT and DM*β* expressing cells. Cells were pre-stained with Calcein Blue and Calcein AM to establish the actual cell ratio in the mixed population (Fig. 4d). The mixed population was then analyzed on the mCPU measuring the pMHC dissociation rate as well as identifying the cell-type using Calcein staining. The mCPU analyzed a total of 376 single-cells in three mCPU runs (Fig. 4e). An initial 90% WT to 10% DM*β* target ratio resulted in an average observed ratio of 88% WT to 12% DM*β* expressing cells as determined by Calcein dye labelling (Fig. 4f). We then sought to identify cells with apparent long half-life pMHC-TCR interactions by setting a half-life threshold of 45s near the lowest observed DM*β* half-life, resulting in a sensitivity of 100%. This threshold level in turn yielded a specificity and accuracy of 96%. The mCPU correctly identified all long half-life DM*β* clones, as well as a small number of WT clones that also exhibited long half-lives (Fig. 4a). With these encouraging results we further increased the input ratio between WT and DM*β* clones by an order of magnitude to 99% to 1%, respectively (Fig. 4g & Supplementary Fig. S12). We let the mCPU run until it detected the first high-affinity cell based on a half-life measurements and were able to identify a DM*β* clone after 45 single-cell measurements.

Enabling downstream processing of selected cells requires identification and recovery of selected cells from the device. We performed an experiment on a mixture of WT and DM*β* TCR expressing cells at a ratio of 77% to 23%, respectively (Fig. 4h). We measured half-lives and specifically routed cells with half-lives above a 45s threshold to a recovery outlet. In the outlet we observed a ratio of 33% WT to 67% DM*β* expressing cells based on Calcein staining, which represents close to an order of magnitude enrichment, based on the input ratio of DM*β* to WT expressing cells of .3 and a post-selection ratio of 2. Cells could be aspirated from the recovery outlet with a pipette and transferred to a multiwell plate and 92% of cells could be successfully transferred with this simple approach (25 cells selected for recovery and 23 cells successfully transferred). We recovered 94% of the selected DM*β* cells. Single-cell recovery and transfer to a multiwell plate was also achieved for the single cell identified and selected in the 99% WT and 1%DM*β* ratio experiment described above (Supplementary Fig. S12).

## Discussion

By enabling complex, time-dependent experiments the mCPU platform fully automates single-cell analysis workflows, and is expected to find applications in areas where precise temporal and spatial control over single-cells is required. For example, promising application areas are basic immunology such as characterizing T and B cells in response to natural infection and applied immunology including cancer immunotherapy where screening large-numbers of T cells and selection of high-affinity candidates is critical for therapeutic applications.

In the case of cancer immunotherapy, rare, high-affinity lymphocytes against tumoral antigens are often found to be scarce (1/1000 or lower). For determining the distribution of TCR affinities in native T-cell populations a throughput of 100 - 1000 would already represent an order of magnitude improvement over recent state-of-the-art measurements [12]. The throughput requirement for screening B-cell libraries strongly depends on the experimental design. With a pre-screen or selection in place, characterizing 1000-10000 cells is most likely sufficient and informative. De novo screening of B-cell or other large libraries requiring 10’000 to 100’000 measurements would be currently not yet achievable with the mCPU.

In this first implementation of the mCPU platform we consistently achieved a throughput of 50 cells per hour and the platform ran robustly for 6 hours. Due to the complete automation of the system it should be possible to setup 2 experiments per day running for 6 hours each, which results in a daily throughput of 600 cells. It is therefore possible to screen up to 3’500 cells in a five day period. The throughput of the mCPU platform is therefore well within range for a number of applications. Reaching higher throughput would require additional improvements. Increasing the robustness of the platform to permit unsupervised run-times of 10 hours (a 2-fold improvement) would increase the daily throughput to 1’000. It may be feasible to reduce the chamber size considerably, allowing more chambers to be included on the mCPU and therefore allowing more cells to be screened per loading round. Likewise, and depending on the application, increasing the number of cells screened per chamber could potentially increase the throughput by an order of magnitude, but will then require two rounds of screening if high enrichment is required.

In biotechnology related areas, the mCPU is capable of characterizing native as well as synthetic receptor libraries, which is useful for T cell, B cell, and de novo engineered cell receptor library characterization and screening. In addition to quantifying cell-surface receptor interactions, the mCPU should also be able to measure fast intracellular signaling events, such as calcium dependent signaling. More broadly, we demonstrate the evolution of microfluidic technology towards a truly autonomous “lab-on-a-chip” platform that performs complex fluid handling operations, data acquisition, data analysis, and decision making in a closed-loop, unsupervised system.

## Acknowledgments

The authors thank Evan Olson for advice on microscopy, data, and image analysis; Manon Blache, Theo Nass, and Sylvain Cam for their help on image analysis, Alexandre Harari for helpful discussions, and Ming Yip for advice on microfluidics and microfabrication. This project received from financial support by the ISREC Foundation, made possible by a donation from the Biltema Foundation, and support by the Ludwig Institute for Cancer Research as well as EPFL.

## Competing interests

The authors declare no competing interests.

## Author contributions

J.S, G.C and S.M conceived the idea to perform single-cell pMHC-TCR dissociation rate measurements. J.S provided the NTAmers, cell clones, performed FACS analysis, and contributed to the methods section. S.J.M. and F.J. invented, designed, and optimized the microfluidic device and methods. F.J. implemented the microfluidic design, wrote software, and performed experiments. F.J. and S.J.M analyzed data and wrote the manuscript.

## Supporting Information

Supporting Information Available: The following files are available free of charge. Detailed materials and methods, extended description of the microfluidic device and the characterization of single-cell dissociation kinetics on-chip.

## Notes

### Competing Interest Statement

The authors have declared no competing interest.

