## Supplementary Information for "High-throughput single-cell TCR - pMHC dissociation rate measurements performed by an autonomous microfluidic cellular processing unit"

### Supporting Information

Materials and Methods

Supplementary Figures

### Materials and methods

#### Cell lines and culture

The SupT1 cell lines used in this study were developed previously as described here [1]. In this project two clones were used: WT (with the WT TCR sequence) and DM$\beta$ with increased affinity compared to WT, within physiological range for cancer neoantigens [2]. Cells were cultured in DMEM media supplemented with L-Glutamine, 10% FBS and 1% Penicillin-Streptomycin (Pen/Strep, Gibco). Cells were passaged once to twice per week at a ratio of 1:20 for 12 maximum passages.

#### NTAmers production & staining

NTAmers were synthetized by the Peptide and Tetramer Core Facility of the Department of Oncology UNIL/CHUV as described previously [3]. Briefly, NTAmers are composed of Streptavidin-Phycoerythrin (SA-PE, Invitrogen) complexed with biotinylated peptides carrying four $Ni2^{+}$-nitrilotriacetic acid (NTA4) moieties and non-covalently bound to His-tagged HLA-A*0201 monomers containing an AF647-labeled $\beta$2m. Monomers were obtained by refolding of the HLA-A*0201 heavy chain in the presence of AF647-labeled $\beta$2m containing the S88C mutation with AF647-maleimide (GE Healthcare) and the analog NY-ESO-$1^{157\text{-}165}$ [SLLMWITQA] tumor antigenic peptide. After purification on a Superdex S75 column, pMHC monomers were mixed at a 10-fold ratio with SA-PE-NTA4 in the presence of $Ni2^{+}$, aliquoted and kept at -80°C.

For staining, generally $5.10^5$ cells were spun down (350g, 5min, 4°C) and re-suspended in 100$\mu$L of buffer (PBS with 4% BSA, 1% Pluronic F-127 (Sigma-Aldrich), 2% Pluronic F-68 (Gibco) and 2% Pen/Strep). Cells were tagged with Calcein (Invitrogen) or Calcein Blue (Invitrogen) for live-cell tagging at a concentration of 5$\mu$M with an incubation of 30min in the dark at 4°C. Following addition of 1mL of buffer, cells were spun down (350 rcf, 5min at 4°C), re-suspend in 100$\mu$L of buffer, and 5$\mu$L of pMHC multimers was added with an incubation at 4°C for 30-45min in the dark. Following incubation, 300$\mu$L of buffer with 40$\mu$L of OptiPrep$^{TM}$ was added to reduce cell sedimentation. The final solution was loaded in a 1.5mL glass vial (SureStop vials, Chromacol, Thermo Scientific) containing a micro magnetic stir bar (PTFE, 2x7mm, Fisherbrand), which was actuated by a micro-stirrer (low speed, Fisherbrand). The glass vials for cells, imidazole-rich buffer and washing buffers were kept on ice and placed on top of the magnetic stirrer (Supplementary

Fig. S1). The chip and microscope stage were enclosed in an environmental chamber kept at 14°C.

##### Dissociation rate measurements by flow cytometry

To validate the single-cell off-rates measurements performed on-chip, SupT1 cells (WT and DM$\beta$) were analyzed for dissociation measurements by flow cytometry as described previously [2]. Briefly, 200'000 cells were incubated for 40 minutes at 4°C with specific NTAmers (HLA-A*0201/ NY-ESO-1$^{157\text{-}165}$) in 50$\mu$L FACS buffer (PBS supplemented with 0.5% BSA, 15 mmol/L HEPES, and 0.02% $NaN_3$). After washing, cells were resuspended in 500$\mu$L FACS buffer at 15°C and cell surface-associated mean fluorescence was measured under constant temperature using a thermostat device (15°C) on a SORP-LSRII flow cytometer(BD Biosciences) following gating on living cells. Multimeric PE-$NTA_4$ scaffold and Alexa647-pMHC monomer fluorescence were measured for 30s (baseline). Imidazole was added at 30sec (100 mmol/L) and PE and Alexa647 fluorescence recorded for 10min. Duplicates were used for each cell clones. Data was processed using the FlowJo software (v9.6, Tree Star, Inc.). After gating on living cells, PE or Alexa647 mean fluorescence intensity was derived using the kinetic module of the FlowJo software. Geometric mean fluorescence intensity over time was then plotted using GraphPad Prism software and monomeric TCR-pMHC dissociation kinetics calculated using a one-phase exponential decay equation.

##### Design and fabrication of the microfluidic device

The device design includes four fluid inputs (with built-in filters) and five fluid outputs. The device features 8 independent chambers that are roughly 150 by 100$\mu$m in size. Heights of 10 and 15$\mu$m (chambers and flow channel, respectively) were chosen to accommodate the typical cell diameter of lymphocytes (around 12$\mu$m). The chambers can be isolated using micromechanical valves (Supplementary Fig. S2). Each chamber can be specifically addressed from the bottom buffer inlet via a multiplexer (*e.g* Chamber #2 is addressed individually in Supplementary Fig. S2). To reduce shear stress and prevent cell loss during buffer exchange and wash steps, each chamber features small sieves (1.5$\mu$m high) to retain cells during fluid exchange operations and two bypass channels to reduce flow velocity in the chamber area. Finally, a partially-closing valve was installed before and after the chamber area. This valve does not fully close when activated, allowing fluid flow while increasing flow resistance.

The microfluidic device was fabricated by standard multilayer soft lithography [4]. The microfluidic device was designed in AutoCAD (Autodesk, California). Molds for the control and flow layers were fabricated on two separate wafers by standard photo-lithographic techniques to obtain microfluidic channels with desired heights. For the control layer, a silicon wafer was primed by oxygen plasma treatment for 7 minutes (TePla 300). SU-8 photoresist (GM 1070, Gersteltec Sarl) was spin-coated to obtain a height of 30$\mu$m. After relaxation (30min) and soft bake (3000s ramp to 130°C, baking for 300s at 130°C and ramping down to 30°C for 3000s), the wafer was exposed under a chrome mask for 16s (365 nm wavelength, 20 mW/cm$^2$ light intensity, Süss MA6 Gen3 mask aligner) followed by a post-exposure bake (2400s ramp to 90°C, baking at 90°C for 2400s and ramping down to 30°C in 2700s). Development of the wafer was performed with propylene glycol monomethyl ether acetate (around 3min) followed by incubation with isopropyl alcohol (2min) to stop the reaction and finally by a hard bake (130°C for 2 hours). For the flow layer, a similar procedure was performed on a separate wafer for the first photoresist layer (sieves, 1.5$\mu$m height, SU-8 GM 1040, 3.2s illumination). The same wafer was used for patterning of the second layer (chamber area, 10$\mu$m height, SU-8 GM 1060, 8.3s illumination). After SU-8 patterning, the wafer was vapor treated with hexamethyldisilazane (HMDS) and AZ 9260 (Microchemical GmbH) was spin-coated to a height of 15$\mu$m on a Süss ACS200 GEN3. The wafer was then exposed under a mask for a total of 40 seconds (Süss MA6Gen3). AZ 9260 photoresist (Microchemical GmbH) was spin-coated for a height of 15$\mu$m and exposed for a total of 40 seconds. The AZ 9260 photoresist was annealed by ramping the temperature to 135°C for 2 hours to generate a rounded profile.

For microfluidic chip fabrication, each wafer was treated with trimethylchlorosilane (TMCS) for silanization by vapor deposition in a desiccator for at least 1 hour. For the control layer PDMS (5:1 ratio elastomer to crosslinker) was prepared, poured over the wafer and degassed in a desiccator for 30min. The wafer was then baked in the oven at 80°C for 25 minutes. Following baking, the PDMS chip was detached by hand from the wafer and punched using a stainless steel punch (OD: 0.65 mm, ID: 0.35 mm, length: 8 mm; Unimed). For the flow layer, PDMS with a 20:1 ratio was spin coated at 1450 rpm (ramping up for 20s, spinning for 35s and ramp down of 20s) onto the wafer and left to sit 15 min before baking for 20 minutes. The layers were then aligned by hand and baked for an additional 90 minutes. The bonded chip was then punched again to create the flow layer interconnects and two output holes were punched with a 2.5cm (OD) manual puncher (Harris Uni-Core). The final chip was then bonded to a glass cover slip (Thickness No. 1, borosilicate glass, VWR Int.) cleaned using Scotch tape. The cover slip was taped to a glass slide for support. Bonding was performed by oxygen plasma treatment of both the PDMS chip and glass cover slip (Femto plasma oven, duration of 20s at 100% power, flow rate of 25 sscm and 0.1 bar).

#### Device setup

Pressurization of the control lines on the microfluidic chip was controlled with pneumatic solenoid valve (Pneumadyne manifolds), pressure levels were set with a pressure regulator (Type 10, 2–60 psi, Marsh Bellofram) and a digital gauge (digital manometer, PDC-102N2BFA, Kobold). A USB relay board (USBRELAY32, Numato Lab) was used to control the electric manifold actuation. Control lines were filled with purified, filtered water (Milli-Q water filtered with 0.45$\mu$m) and slowly pressurized to 125 kPA. The flow layer was primed with a buffer (PBS with 4% BSA, 1% Pluronic F-127, 2% Pluronic F-68 and 2% Pen/Strep). Tygon tubing (OD 0.0600, ID 0.0200; Cole-Parmer) with stainless steel connecting pins (0.35mm x 8mm, AISI304, Unimed) was used for control lines and buffer solutions. For cell loading, a combination of connecting pin, Tygon tubing and a narrower tubing was used (PEEK, 1/32" OD, .005" ID, Vici) to prevent cell sedimentation in the tube during the experiment. Cells were loaded at desired densities (usually $5.10^5$ cells in 400$\mu$L) in NTAmer-rich buffer (see above). After initial loading and user-based adjustments (chamber position saving, camera and experimental settings), the custom MATLAB GUI conducted the entirety of the experimental cycle described. Valve actuation sequences are defined for key steps in Supplementary Figure S2. Cell recovery was performed from the larger outlet holes (2.5mm), allowing direct pipetting of cells. Recovered cells were transferred into a flat bottom, square-well 384-well plate (Nunclon Delta Surface, Thermo Scientific) and visualized on a microscope.

#### Data acquisition & analysis

Solenoid valves, microscope, and camera were controlled by the custom Matlab GUI. The chip and microscope stage were enclosed in an environmental chamber kept at 14°C. Imaging of the microfluidics device was performed on a Nikon Ti-E Eclipse automated microscope with an epi-fluorescence illuminator (C-HGFI Intensilight, Nikon). Images were acquired with an Ixon DU-888 camera (EMCCD camera, Andor Technology), using a 60x oil objective (CFI Plan Apo Lambda 60X Oil, Nikon). Time-lapses of fluorescent signals were recorded for two fluorescent labels. The first fluorescent tag uses Alexa647 probes linked to the pMHC monomers. The second fluorescent tag is a PE probe linked to the core unit maintaining the pMHC monomers in a multimeric form. After addition of imidazole-rich buffer (200 mM of imidazole in PBS with 4% BSA, 1% Pluronic F-127, 2% Pluronic F-68 and 2% Pen/Strep), the His-Tag links between the PE core and the Alexa647-fluorescent pMHC monomers is disrupted by competitive inhibition and only the monomeric signal is kept and recorded. This dual use of fluorescent tags allows to determine the time component of the switch from multimeric to monomeric form. Brightfield images (exposure time: 50ms, 50 gain, EM gain x5.1, temperature at -70°C) as well as fluorescent images for both the Alexa647 fluorophores (Nikon filter cube: Cy5, HC 620/60, HC 700/75, BS 660; exposure time: 250ms, 250 gain, EM gain x5.1, temperature at -70°C) and PE fluorophores (Nikon filter cube: Cy3 HC 535/40, HC 590/40, BS 565; exposure time: 250ms, 250 gain, EM gain x5.1, temperature at -70°C) were recorded during time-lapses for a total of 60 to 90 seconds with a fixed interval of 3s for each channel for a total of 20 to 30 frames acquired per channel. Fluorescent images were corrected by dark frame subtraction and flat-field correction. Timeseries images were analyzed using custom MATLAB code. Briefly, cells were identified as ROIs. ROIs were defined using Calcein tag (FITC channel, exposure time: 25ms, 25 gain, EM gain x5.1, temperature at -70°C), Calcein Blue (DAPI channel, exposure time: 125ms, 125 gain, EM gain x5.1, temperature at -70°C) or brightfield images (via trained CNN or ACF object detectors) around cells of interest and around background regions (Supplementary Fig. S3). The mean intensity of the ROIs was calculated over the different frames, the background mean intensity was subtracted from the mean signal intensity and the signal over time was plotted and fitted to a one-phase exponential decay corrected for bleaching. Bleaching rates were established for both fluorophores by performing flow exchange with buffer lacking imidazole. This allowed to extract bleaching rates for Alexa647 and PE of 245s and 123s respectively, Supplementary Fig. S4. The bleaching rate of Alexa647 was used to compensate bleaching during experiments by adapting dissociation rates accordingly ($A * e^{(\text{-}(\mathrm{k}+\mathrm{k}_{\mathrm{bleaching}})^{*}\mathrm{t})}+B$).

After fitting, key parameters were extracted for event filtering. The key parameters included goodness of fit ($R^2$) keeping events with a minimum of 0.95 to ensure proper fitting of the data. Other key parameters were initial intensity (A, maximum level of 45'000 RFUs) and background level (B, maximum level of 15'000 RFUs). Finally, a drop in PE signal (difference between initial and final intensity) higher than 32% . In addition of those key parameters, the half-lives of dissociation were extracted after filtering and fitting (Supplementary Figures S6, S9 & S10). The extracted half-lives were used to trigger the decision on cell selection for output and recovery.

#### Code availability

The complete MATLAB code is included in the **Supplementary Software** zip file.

### Supplementary Figures

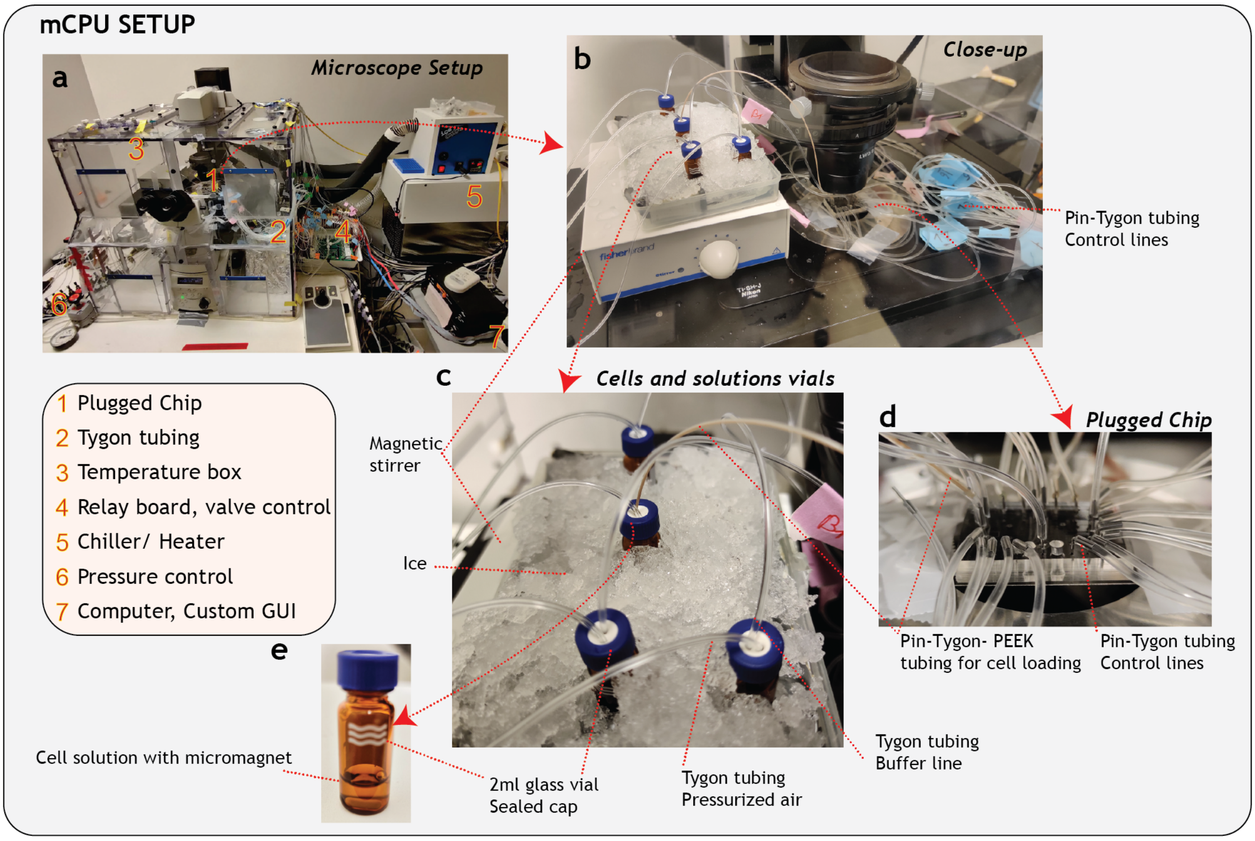


Figure S1: **mCPU setup**. **a** Microscope setup with connected mCPU, temperature control, pneumatic valve and computer control. **b** Close-up view of the mCPU device and connected buffer and cell vials. **c** Vials with buffers and cells were kept on ice to prevent temperatures to reach above 15°C. Tygon and PEEK tubing is used to connect the vials to the chip. **d** Image of the mCPU with larger punched outlets for Outputs 1 and 2 for cell recovery. **e** Images of a glass vial used for buffers and cells. The cell vial includes a micromagnet for stirring.

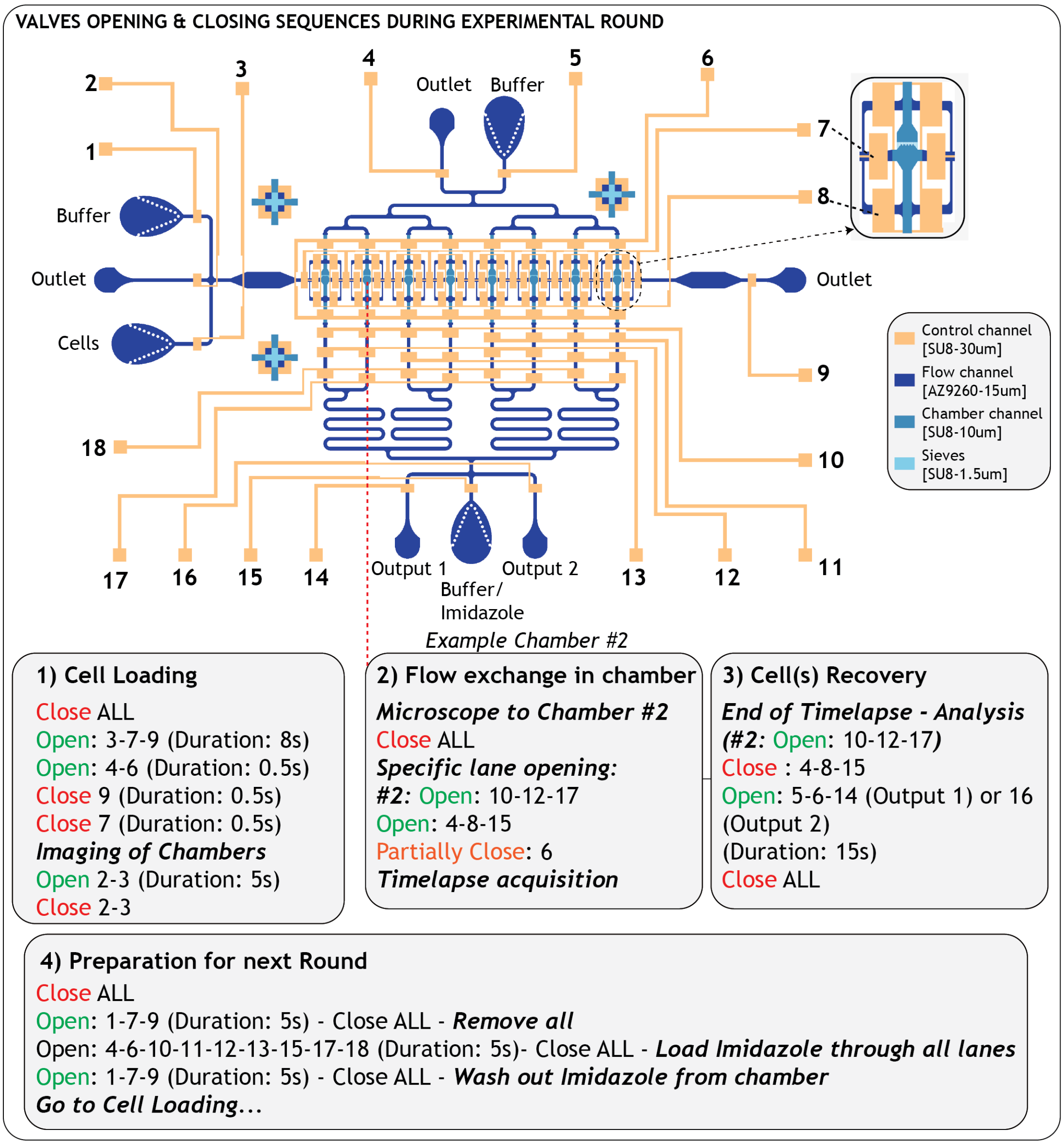


Figure S2: **Pneumatic valve actuation states and sequences during the various experimental round**. For each major step of the experimental cycle, valves are actuated in a specific sequence and for a given time. Each chamber is addressed independently by a multiplexer for flow exchange and cell recovery (here, chamber #2 is addressed).

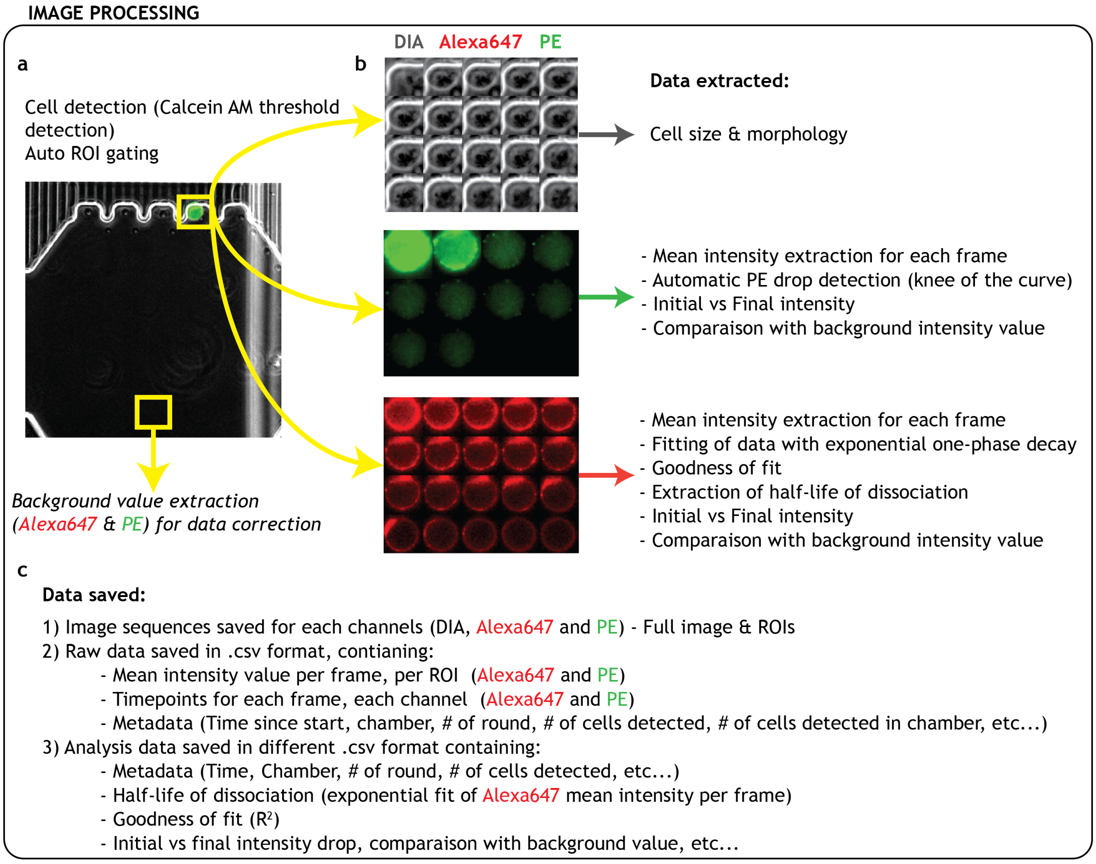


Figure S3: **"Real-time" image processing**. **a** Automated cell detection is performed either via brightfield detection (CNN/ACF) or via fluorescent live-cell tags (*e.g.* Calcein AM). Automatic ROIs are created for each detected cell and one background ROI at the bottom of the chamber for background subtraction. **b** After ROI creation, each frame is automatically analyzed and data is extracted for each channel. Brightfield channel provides cell size and morphology information. The mean intensity of the PE channel (multimeric core of the *NTAmers* technology) is analyzed to automatically detect the sudden drop in signal, indicating a switch from multimeric to monomeric states of the NTAmer (code automatically identifies knee of curve). After the monomeric switch timepoint is identified, the analysis of the mean intensity of the Alexa647 channel is cropped accordingly and the data is fitted with an exponential one-phase decay to extract the dissociation half-life of the cell analyzed. Additional data such as goodness of fit ($R^2$), initial intensities, background level intensity, *etc...*, are also recorded. **c** All images and data generated are saved in either .tiff or .csv format.

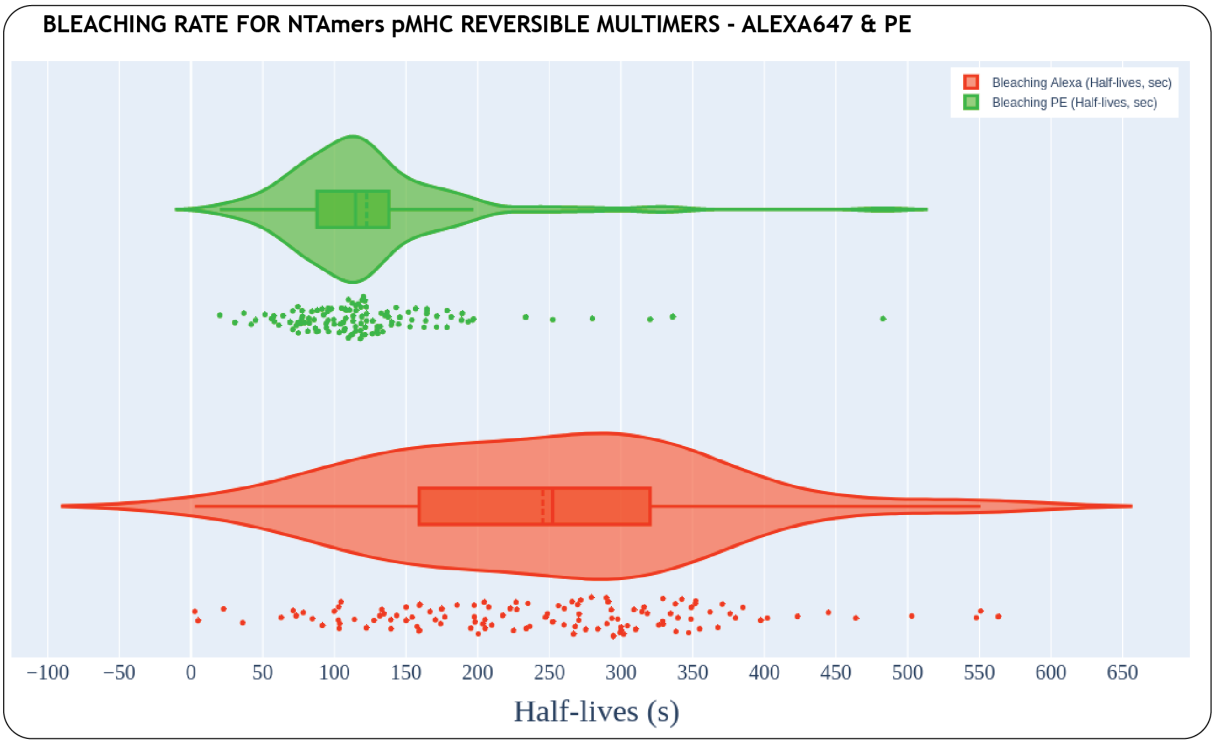


Figure S4: **Bleaching rates for NTAmer Alexa647 and PE fluorophores**. Bleaching rates were established on the mCPU by performing flow exchange with buffer without imidazole with same parameters (camera, time-lapse duration, *etc...*) as normal experimental conditions. The obtained bleaching rate for Alexa647 was used for bleaching correction during fitting ($A * e^{(\text{-}(\text{k}+\text{k}_{\text{bleaching}})^{*}\text{t})} + B$).

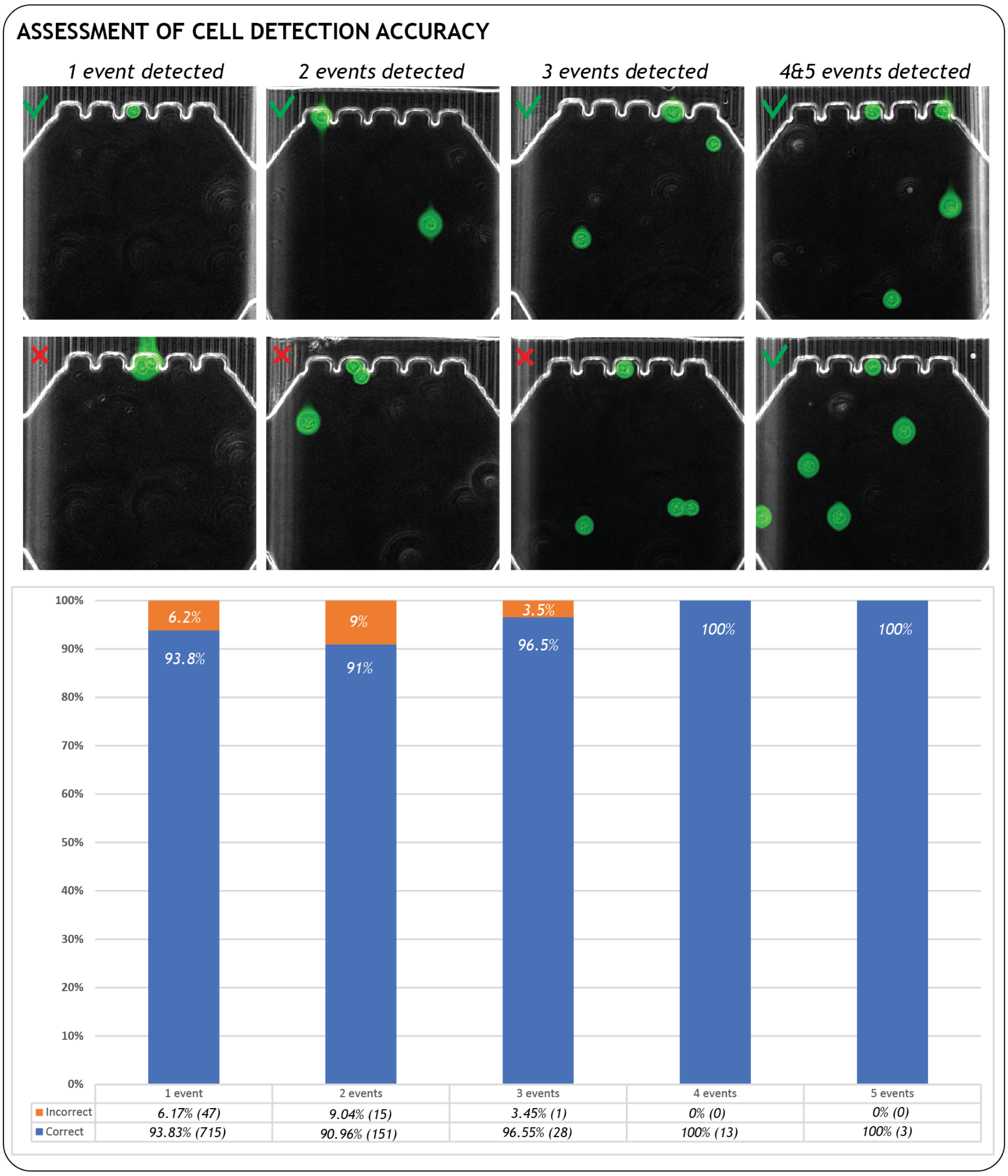


Figure S5: **Assessment of cell detection accuracy** Saved images with Calcein AM hits were analyzed to detect errors in cell detection in comparison with saved metadata. The recurrence of errors in calculated events per chambers (*e.g.* 2 cells touching triggering a 1-cell detection event) was assessed over the different experiments performed.

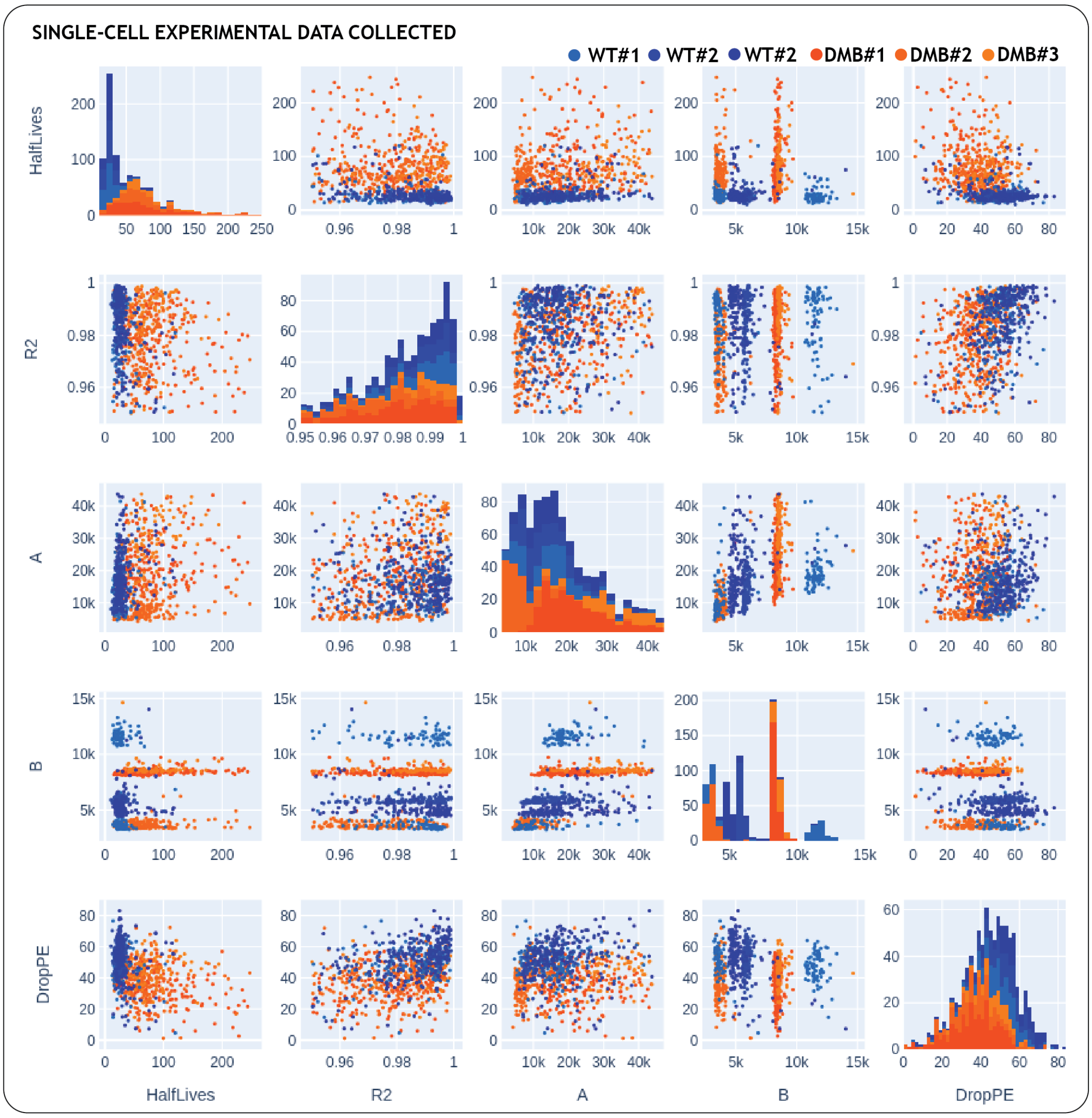


Figure S6: **Single-cell experimental data collected** Scatter plot of single-cell events collected during 6 experiments (3 on WT clones, 3 on DM$\beta$ clones). Data points are shown for events that passed the filter criteria. Key parameters extracted and used for fitting and filtering are goodness of fit ($R^2$), A (initial Alexa intensity), B (background level), and drop in PE, which represents the difference between initial and final PE intensity levels.

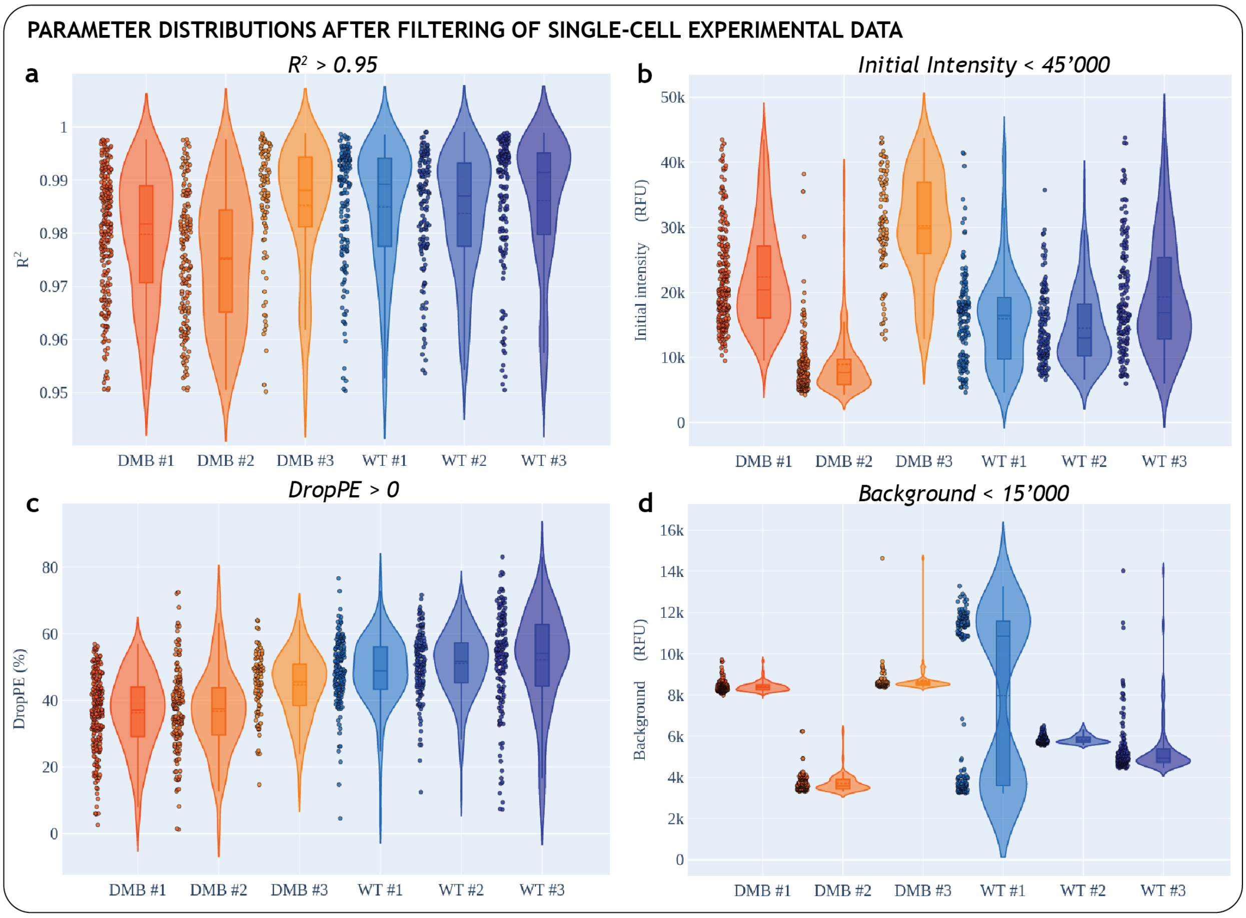


Figure S7: **Parameter distributions after filtering of single-cell experimental data a** Goodness of fit ($R^2$ distribution post-filtering for the different experimental replicates. **b** Initial intensity (A) distribution. **c** Distribution of drop in PE, the difference between initial and final intensity in PE. **d** Background intensity (B) distribution.

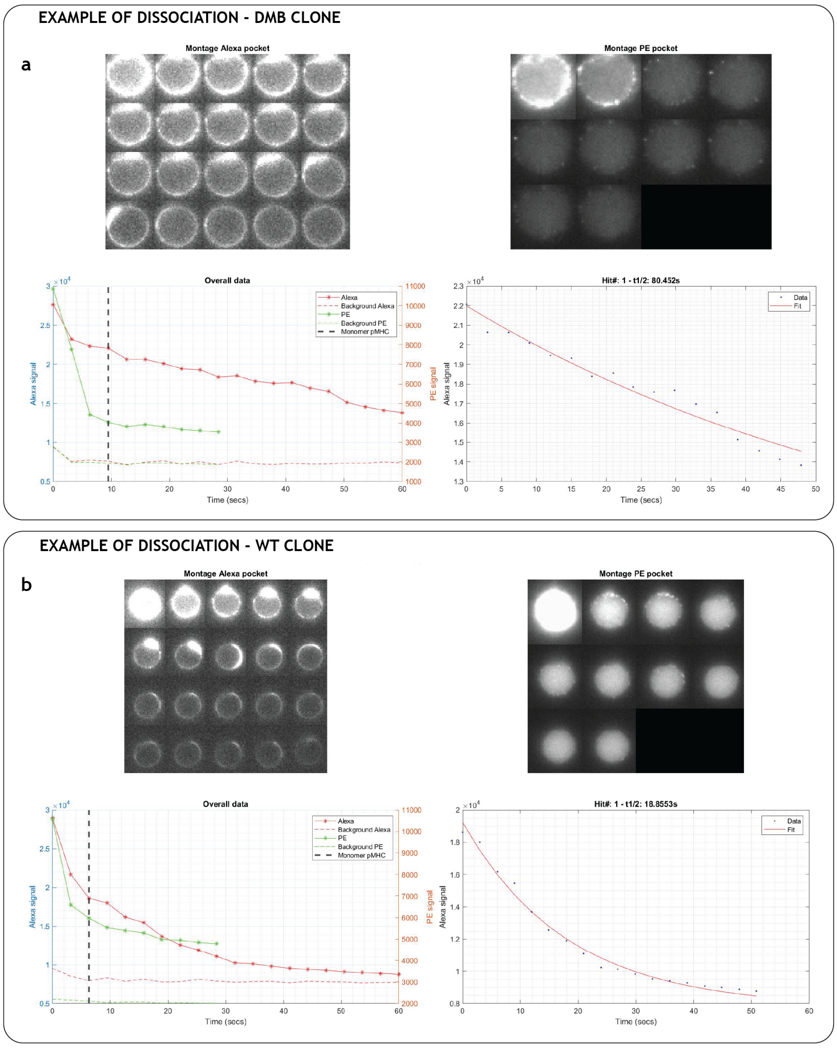


Figure S8: **Examples of automatically-generated data of single-cell pMHC-TCR dissociation measurements**. Data are automatically generated and plotted during experimental cycle for a single event. **a** Example of dissociation kinetics of a single DM$\beta$ cell with image timeseries of both Alexa647 and PE signals and overall mean intensity plots (bottom left plot). The sharp decrease in PE signal is automatically detected and is shown by the vertical black dotted line. The mean intensity of the Alexa647 is cropped accordingly and then fitted with a one-phase exponential decay to extract half-life of dissociation (here, around 80 seconds). **b** Similar example for a single WT cell ($t_{1/2}$ = 19 seconds).

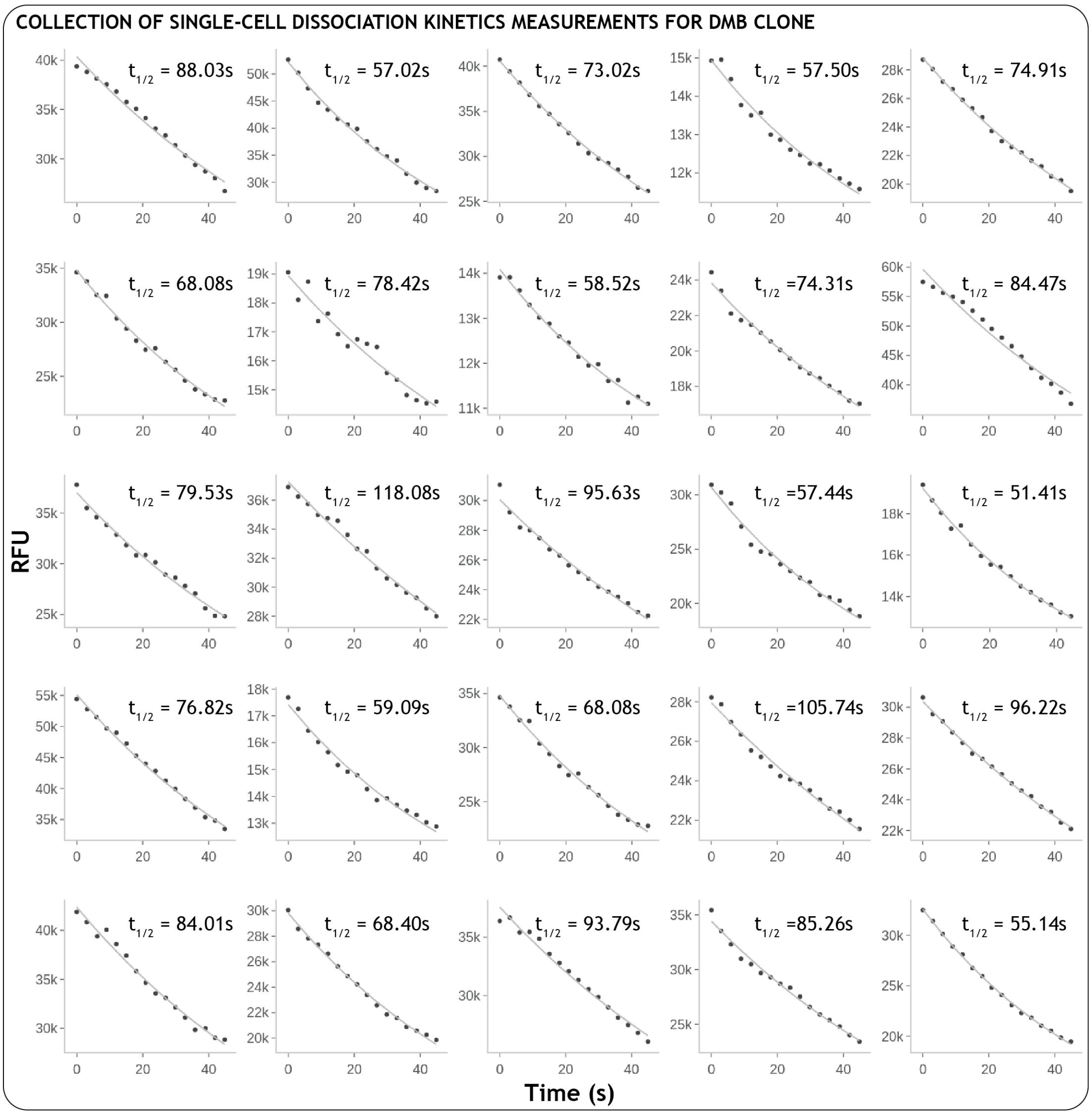


Figure S9: **Examples of single-cell dissociation kinetics measurements for DM$\beta$ cells**. Random selection of 25 filtered events during DM$\beta$ dissociation measurements. Data was automatically cropped to the time when the switch to monomer (PE drop) occurred.

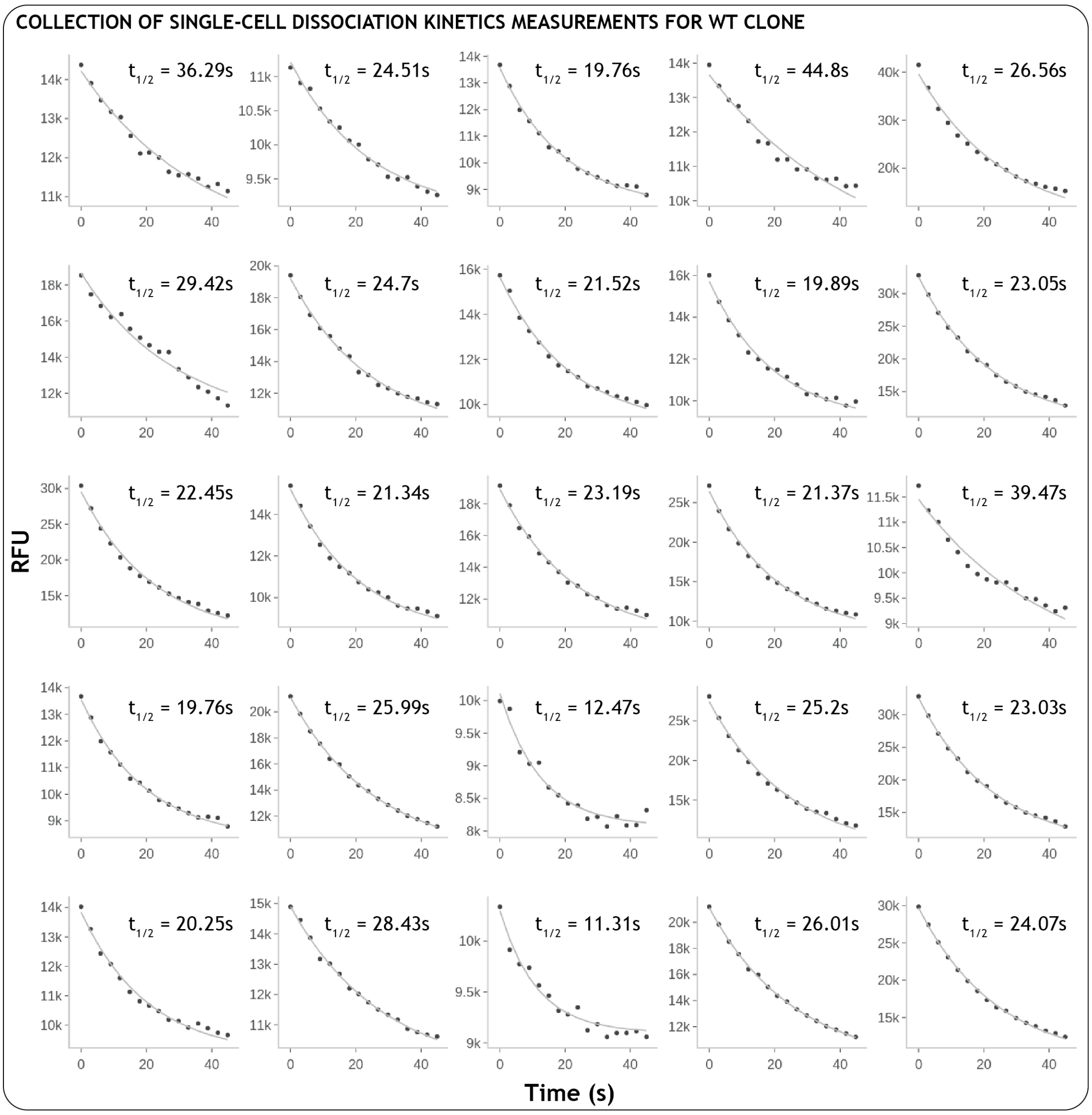


Figure S10: **Examples of single-cell dissociation kinetics measurements for WT cells** Random selection of 25 filtered events during WT dissociation measurements. Data was automatically cropped to the time when the switch to monomer (PE drop) occurred.

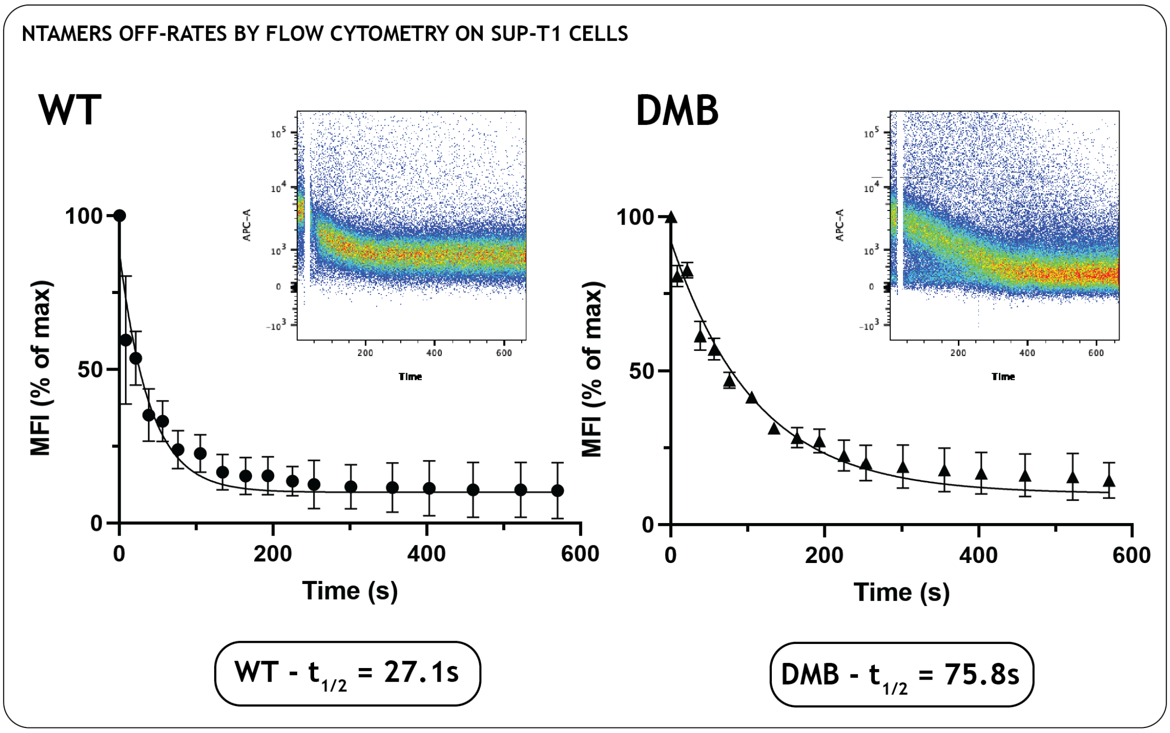


Figure S11: **pMHC-NTAmers dissociation rates determined by flow cytometry**. Sup-T1 cells from same batch as the clones used during single-cell experiments on-chip were analyzed by flow cytometry for off-rates measurements. The extracted half-lives were compared with values obtained on-chip for validation. WT clones showed a half-life of dissociation of 27.1s and DM$\beta$ clones showed an average half-life of 75.8s by flow cytometry.

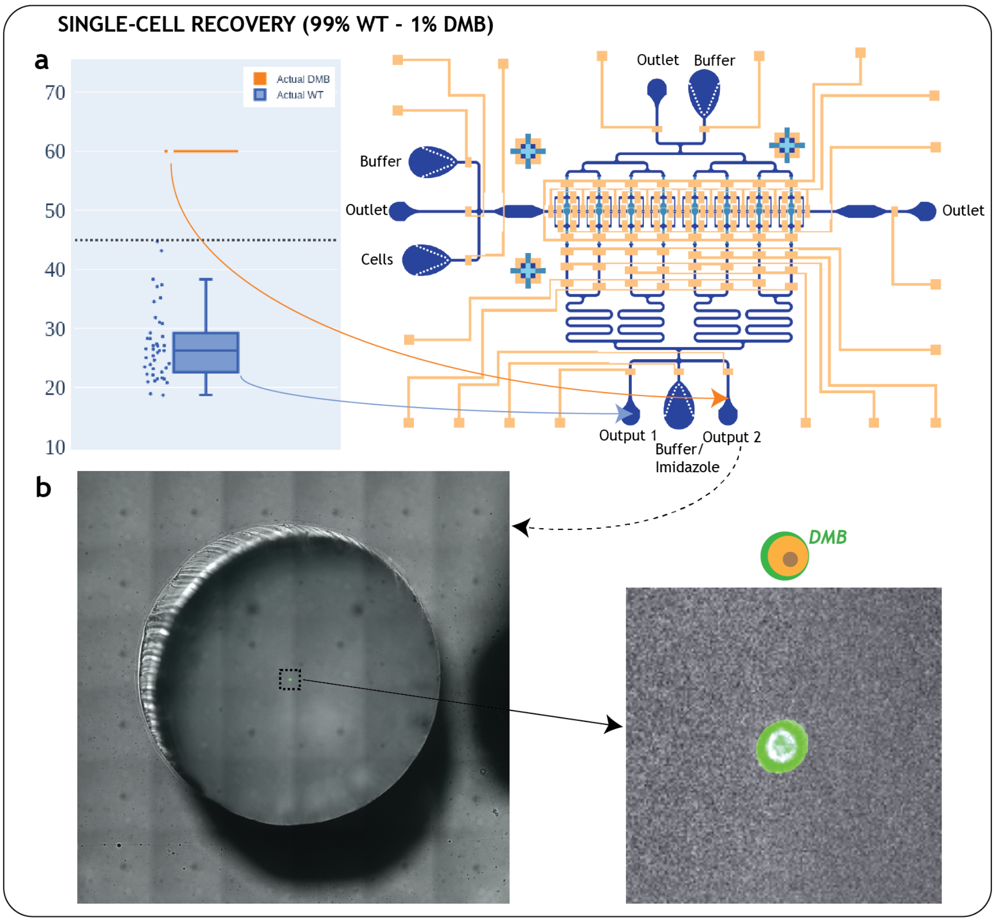


Figure S12: **Single-cell recovery a** A mixed cell population (99% WT - 1%DM$\beta$) was tagged with Calcein Blue and Calcein AM respectively and differentiated by half-life thresholding. The first cell encountered with half-life superior than 45s was pushed to Output 2. **b** Image of single recovered cell. Imaging confirmed the cell to be a DM$\beta$ clone.

[2] Michael Hebeisen, Mathilde Allard, Philippe O. Gannon, Julien Schmidt, Daniel E. Speiser, and Nathalie Rufer. Identifying individual T cell receptors of optimal avidity for tumor antigens. *Frontiers in Immunology*, 6(NOV), 2015.

[3] Julien Schmidt, Philippe Guillaume, Melita Irving, Petra Baumgaertner, Daniel Speiser, and Immanuel F. Luescher. Reversible major histocompatibility complex I-peptide multimers containing Ni 2+-nitrilotriacetic acid peptides and histidine tags improve analysis and sorting of CD8 + T cells. *Journal of Biological Chemistry*, 286(48):41723–41735, 12 2011.

[4] Todd Thorsen, Sebastian J. Maerkl, and Stephen R. Quake. Microfluidic Large-Scale Integration. *Science*, 298(5593):580–4, 10 2002.
